## Supplementary Figures and Figure Legends for "JASPer controls interphase histone H3S10 phosphorylation by chromosomal kinase JIL-1 in *Drosophila*"

Supplementary Figure 1 - Albig et al.

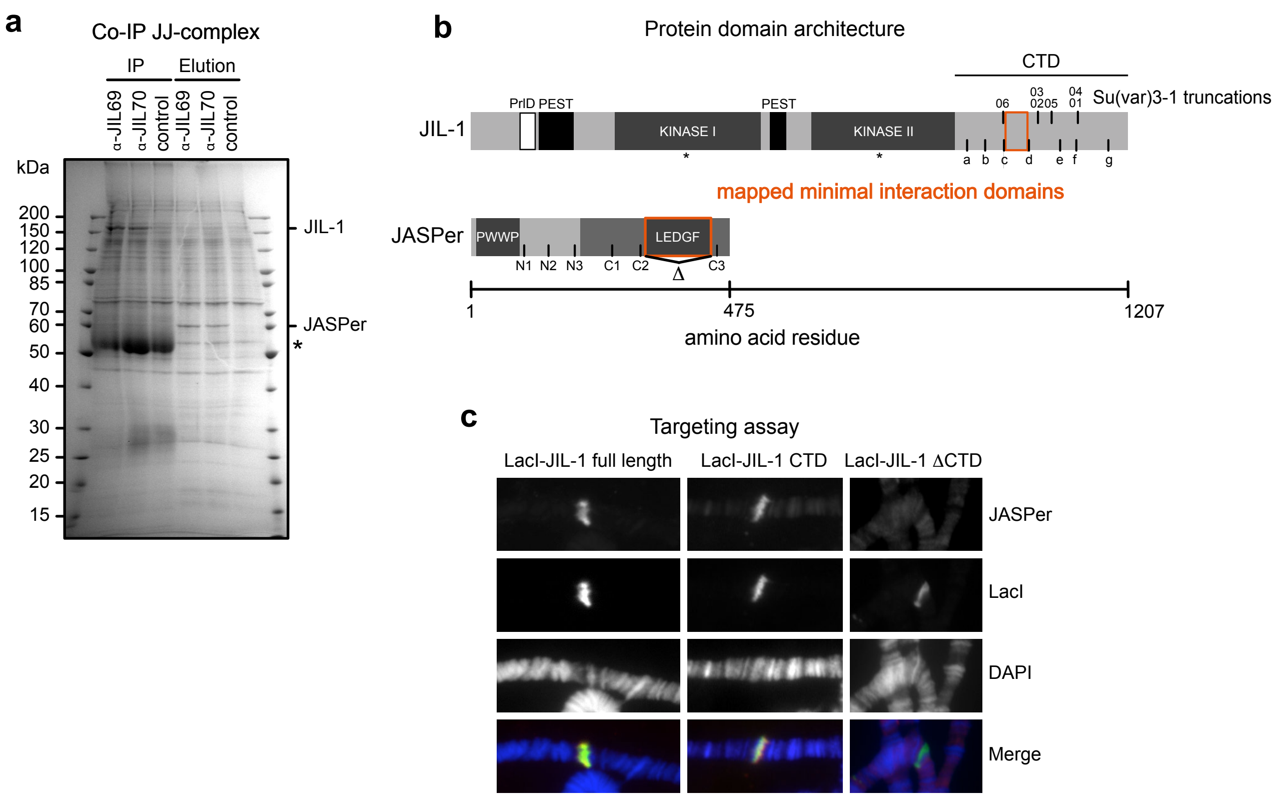

**Supplementary Figure 1. JASPer interacts with the CTD of JIL-1.**

(a) SDS-PAGE with Coomassie staining of JJ-complex co-IP from nuclear embryo extracts. Co-IP was performed with two different $\alpha$-JIL-1 serum and sarkosyl elution fraction is loaded on the right to the IPs. Antibody heavy chain is marked by asterisk.

(b) JIL-1 and JASPer domain architecture drawn to scale as in Figure 1c. In JIL-1, PEST domains are highlighted in black, kinase domains in dark grey and a predicted prion like domain in white. Su(var)3-1 truncations are marked by lines on upper site and deletion mutants on lower site. In JASPer, PWWP and LEDGF domains are highlighted in dark grey and conserved region in intermediate grey. Deletion mutants are marked by lines at lower site, with $\Delta$ denoting JASPer-$\Delta$LEDGF.

(c) Immunofluorescence microscopy of LacO-LacI targeting assay on polytene chromosome squashes from L3 larvae homozygous for the *LacO* repeat line P11.3 using LacI-JIL-1 full length, LacI-JIL-1 CTD and LacI-JIL-1 $\Delta$CTD. From top to bottom, staining for JASPer, JIL-1, DNA and merged images are shown.

Supplementary Figure 2 - Albig et al.

a

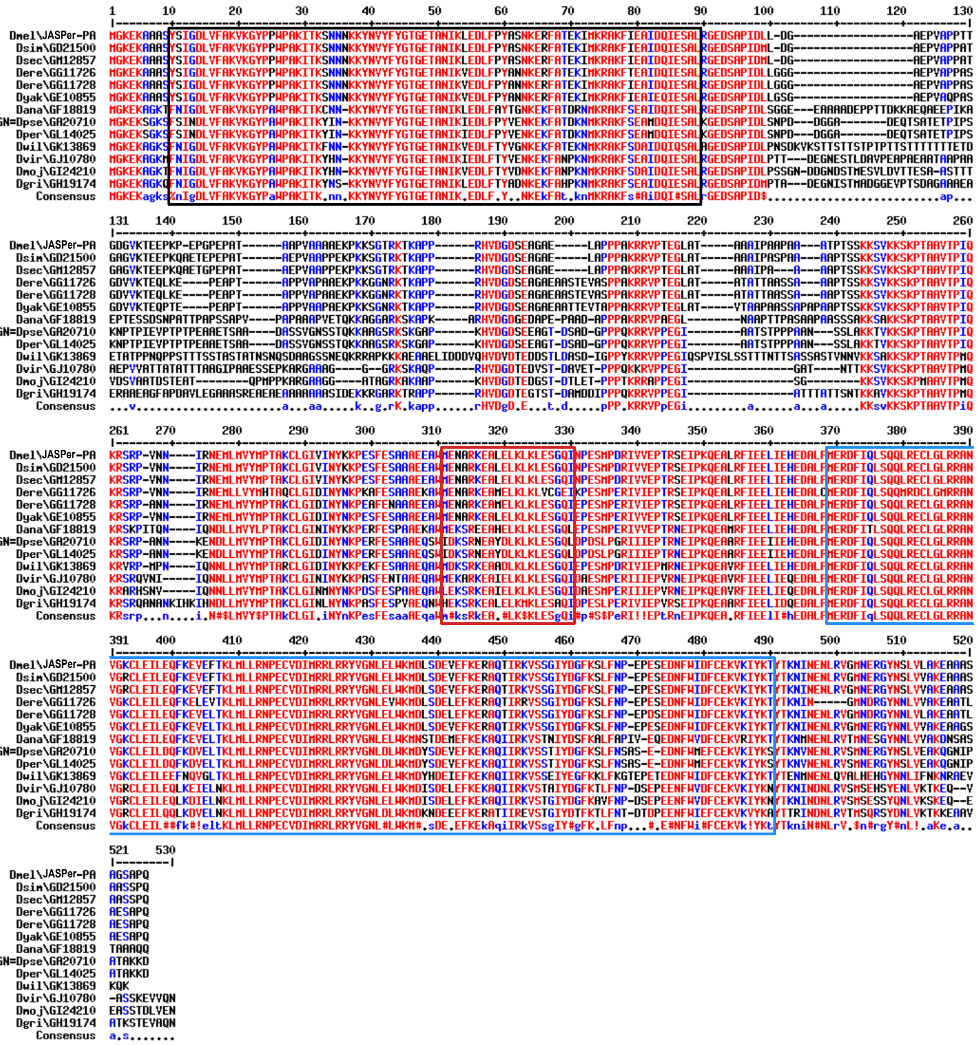

b

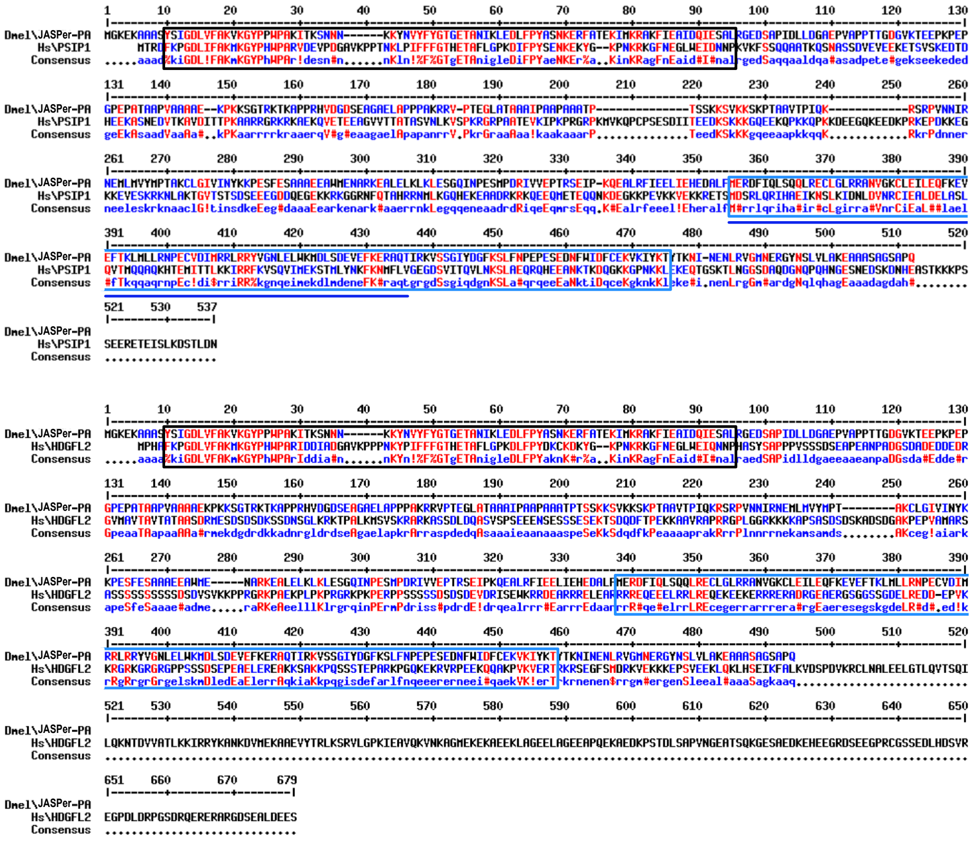

**Supplementary Figure 2. Conservation of JASPer from flies to human.**

(a) Alignment of JASPer homologs from 11 *Drosophila* species (Dmel: *D. melanogaster* JASPer-PA; Dsim: *D. simulans* GD21500; Dsec: *D. sechellia* GM12857; Dere: *D. erecta* GG11726 and GG11728; Dyak: *D. yakuba* GE10855; Dana: *D. ananassae* GF18819; Dpse: *D. pseudoobscura* GA20710; *D. persimilis* GL14025; Dwil: *D. willistoni* GK13869; Dvir: *D. virilis* GJ10780; Dmoj: *D. mojavensis* GI24210; Dgri: *D. grimshawi* GH19174) using MultAlin [98]. Residues conserved to more than 90% are marked in red and residues conserved to 50-90% in blue. The PWWP domain is marked by a black rectangle, the predicted coil domain by red and the LEDGF domain by light blue.

(b) Alignment of *D. melanogaster* JASPer with human PSIP1 in upper panel and with human HDGFL2 in lower panel. Residues are conserved to more than 90% are marked in red and residues conserved to 50-90% in blue. The PWWP domain is marked by black rectangle, the LEDGF domain by light blue and the IBD by dark blue line.

### Supplementary Figure 3 - Albig et al.

a

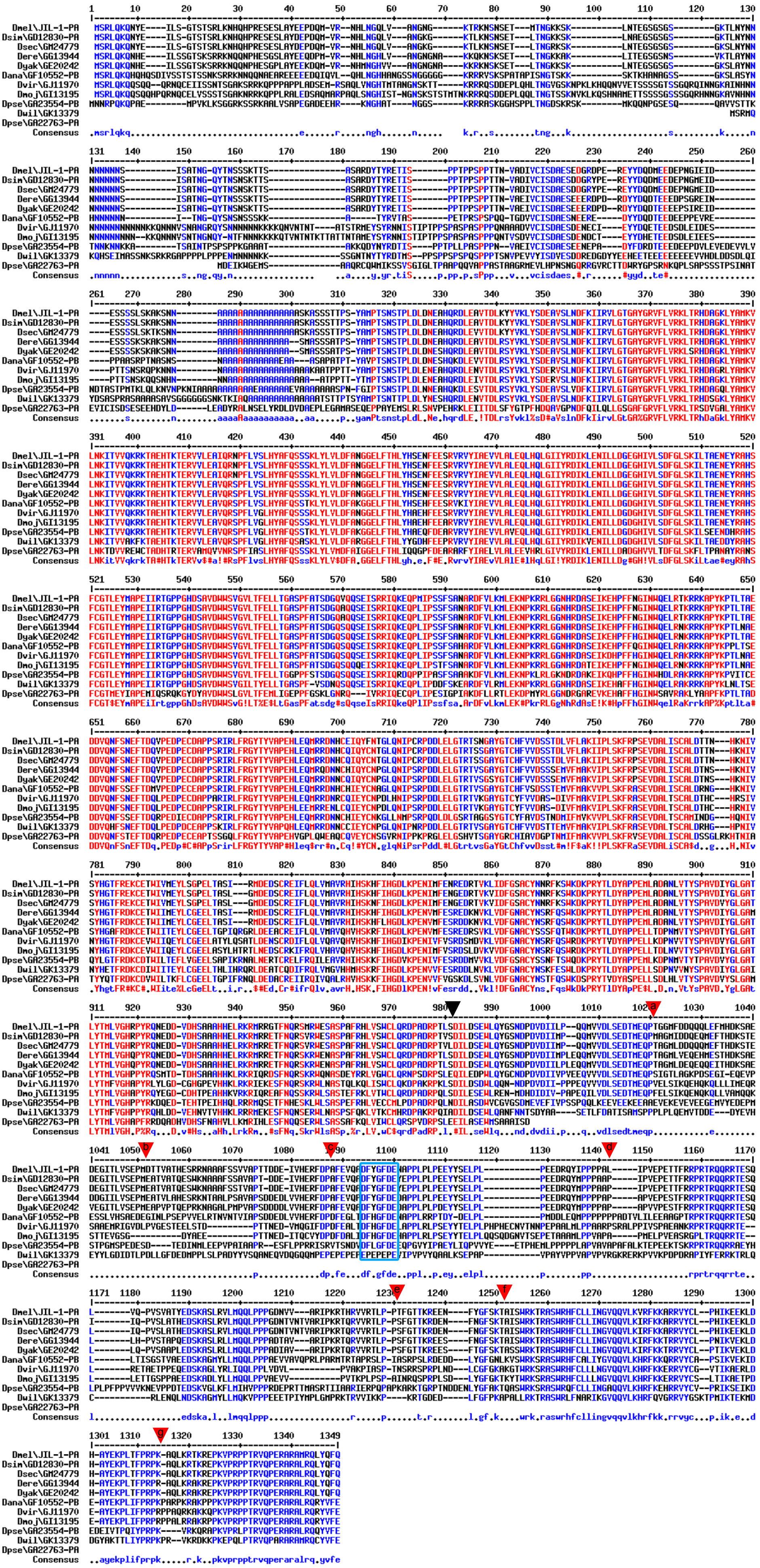

**Supplementary Figure 3. Conservation of JIL-1 in Drosophilae.**

(a) Alignment of JIL-1 homologs from 9 *Drosophila* species (Dmel: *D. melanogaster* JIL-1-PA; Dsim: *D. simulans* GD12830-PA; Dsec: *D. sechellia* GM24779; Dere: *D. erecta* GG13944; Dyak: *D. yakuba* GE20242; Dana: *D. ananassae* GF10552-PB; Dvir: *D. virilis* GJ11970; Dmoj: *D. mojavensis* GI13195; Dpse: *D. pseudoobscura* GA23554-PB and GA22763-PA; Dwil: *D. willistoni* GK13379) using MultAlin [98]. Residues are conserved to more than 90% are marked in red and residues conserved to 50-90% in blue. The IBM is marked by light blue rectangle, the end of kinase domain II by black arrow head and the truncation used for interaction mapping by red arrow heads.

Supplementary Figure 4 - Albig et al.

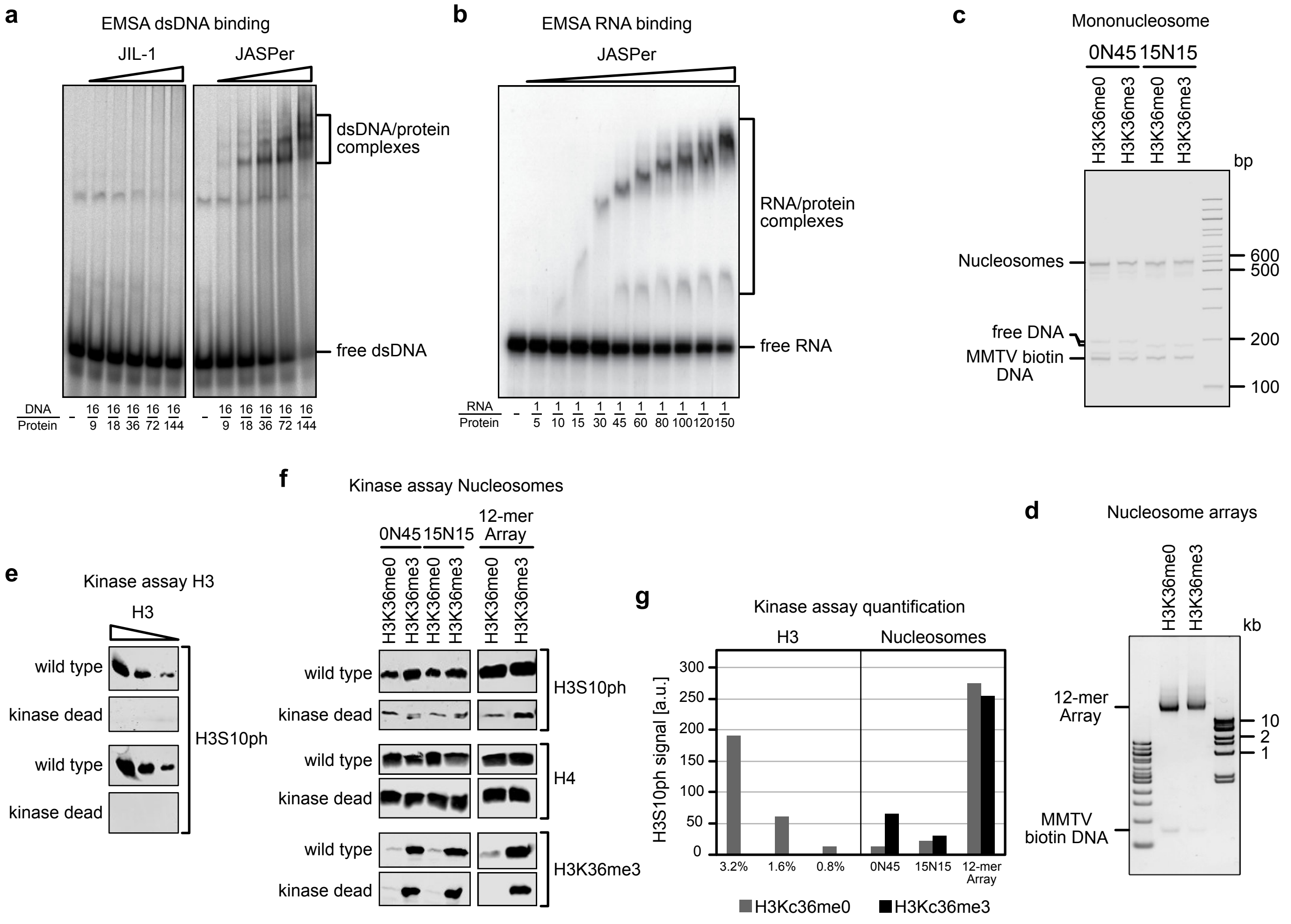

**Supplementary Figure 4. Biochemical characterization of the JJ-complex.**

(a) EMSA with fluorescence detection of recombinant purified JIL-1 and JASPer, left and right panel respectively, binding to dsDNA resolved by native PAGE. Molar ratio of protein to 70 nM of 40 bp Cy5-labeled dsDNA corresponding to the end of the 601 positioning sequence is indicated below.

(b) EMSA with phosphoimaging detection of recombinant purified JASPer binding to RNA resolved by native PAGE. Molar ratio of protein to 2.5 nM 123 bp $^{32}$P-labeled RNA is indicated below.

(c) Agarose gel electrophoresis with SYBR Gold nucleic acid staining of mononucleosomes with unmodified and H3K36me3 modified nucleosomes with 0N45 and 15N15 DNA are loaded next to each other. Molecular weight markers are shown to the right.

(d) Agarose gel electrophoresis with SYBR Gold nucleic acid staining of 12-mer nucleosome array with unmodified and H3K36me3 modified are loaded next to each other. Molecular weight markers are shown to the left and right.

(e) Western blot analysis using $\alpha$-H3S10ph antibody of kinase assay on histone H3 using JJ-complex and kinase dead mutant. We first defined conditions in a radioactive kinase assay to reach a 1:1 molar phosphorylation ratio of isolated H3 with the wild-type JJ-complex, and performed cold kinase assays in the same conditions with the same amount of wild-type JJ-complex and of the mutant JJ-complex containing the kinase dead JIL-1 (D407A and D759A). Using 0.8 – 3.2 % of the 1:1 phosphorylated H3, we confirmed that only the wild type JJ-complex phosphorylates H3S10 and that we can detect little amounts of H3S10ph by western blot.

(f) Western blot analysis using $\alpha$-H3S10ph, $\alpha$-H3K36me3 and $\alpha$-H4 antibodies of kinase assay on mononucleosomes and 12-mer nucleosome array using JJ-complex and kinase dead mutant. Unmodified and H3K36me3 modified nucleosomes are loaded next to each other. We tested two types of mononucleosomes either with symmetric DNA overhang (15N15: 15 bp DNA on each side) or with a protruding DNA overhang only on one side (0N45: 45 bp DNA only on one side), as well as 12-mer nucleosome arrays with or without H3K36me3. For the kinase assay with mononucleosomes and 12-mer nucleosome arrays, we used the same conditions as in (e) but had to load ~10-times more of each reaction to detect similar levels of H3S10ph by western blot. Due to the increased loading, we observed low H3S10ph signals with the mutant JJ-complex with kinase dead JIL-1 and assumed that this is background sticking of the antibody to unmodified H3 and subtracted the values for further analysis shown in (g).

(g) Bar plot of quantification of Western blots presented in (e) and (f). Amount of H3 kinase assay loaded is indicated below.

Supplementary Figure 5 - Albig et al.

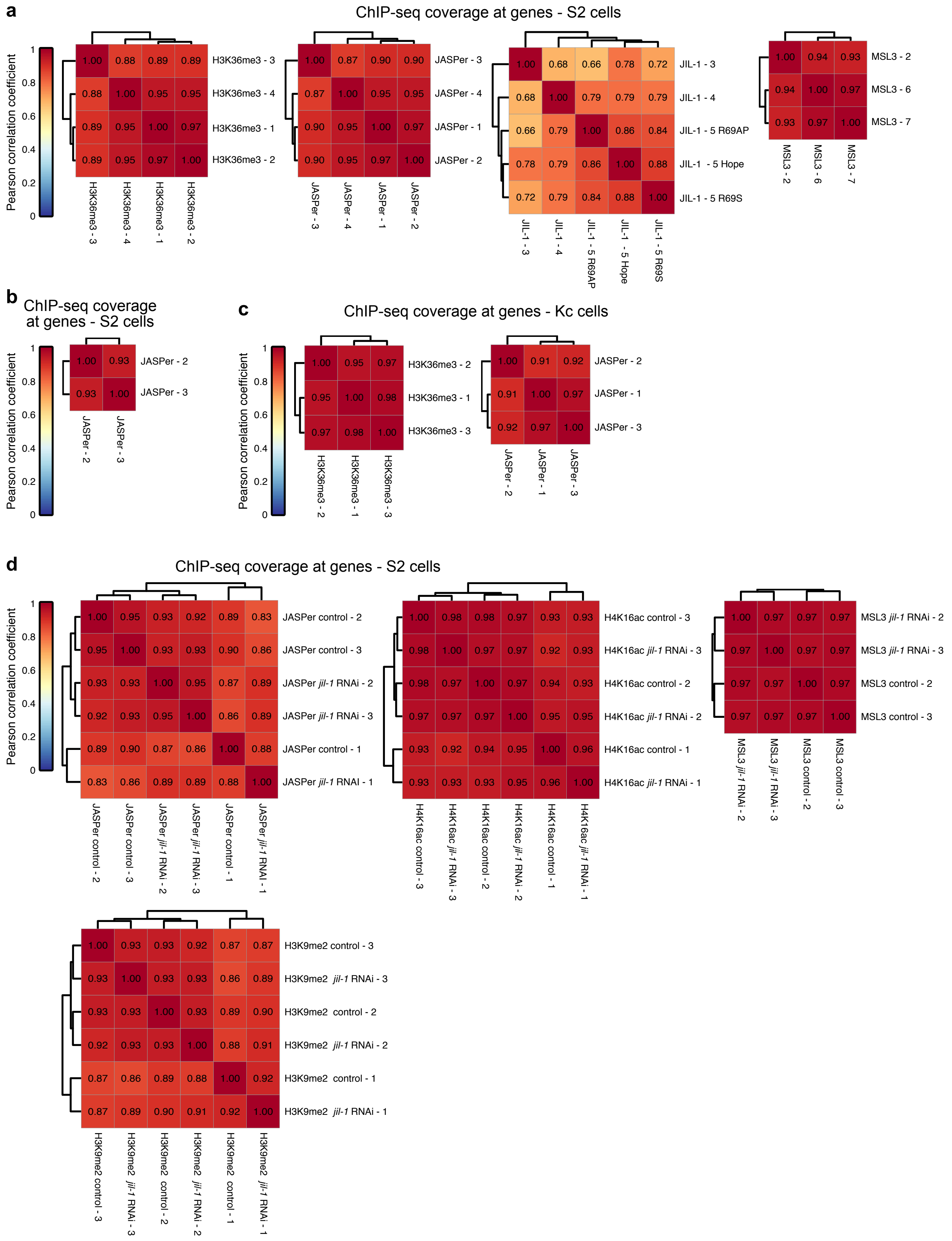

**Supplementary Figure 5. Correlation between biological ChIP-seq replicates at genes.**

(a) Heatmap showing Pearson correlation coefficients between biological replicates of normalized MNase ChIP-seq coverages at genes (n = 17452) in male S2 cells, from left to right for H3K36me3, JASPer, JIL-1 and MSL3.

(b) Heatmap showing Pearson correlation coefficients between biological replicates of normalized JASPer sonication ChIP-seq coverages at genes (n = 17452) in male S2 cells.

(c) Heatmap showing Pearson correlation coefficients between biological replicates of normalized MNase ChIP-seq coverages at genes (n = 17452) in female Kc cells, from left to right for H3K36me3 and JASPer.

(d) Heatmap showing Pearson correlation coefficients between biological replicates of normalized spike-in ChIP-seq coverages at genes (n = 17452) in control male S2 cells and after *jil-1* RNAi treatment, from left to right on top for JASPer, H4K16ac, MSL3 and on bottom H3K9me2.

Supplementary Figure 6 - Albig et al.

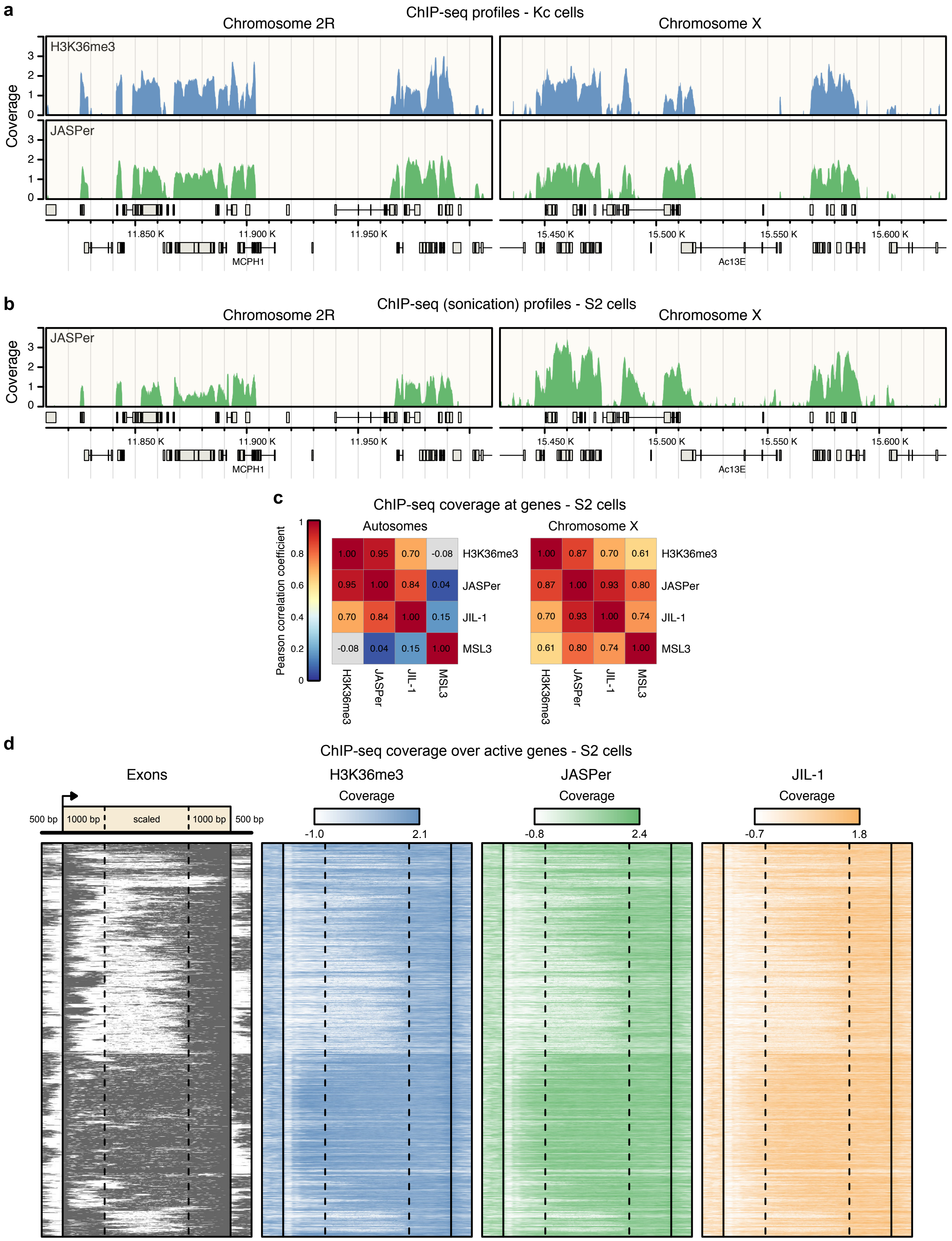

**Supplementary Figure 6. JJ-complex and H3K36me3 are enriched at exons of active genes.**

(a) Genome browser profile showing mean H3K36me3 (upper panel) and JASPer (lower panel) MNase ChIP-seq normalized coverage (n = 3 each) along representative 200 kb windows on chromosome 2R and X in female Kc cells. Gene models are drawn below in grey.

(b) Genome browser profile showing normalized mean JASPer Sonication ChIP-seq coverage (n = 2) along representative 200 kb windows on chromosome 2R and X in male S2 cells. Gene models are drawn below in grey.

(c) Heatmap showing Pearson correlation coefficients between mean MNase ChIP-seq normalized coverages at autosomal genes (left panel, n = 14511) and X chromosomal genes (right panel, n = 2621) in male S2 cells for H3K36me3 (n = 4), JASPer (n = 4), JIL-1 (n = 5) and MSL3 (n = 3).

(d) Heatmap showing exons (left) and mean H3K36me3 (second left, n = 4), JASPer (second right, n = 4) and JIL-1 MNase ChIP-seq normalized coverages (right, n = 5) at active (tpm > 1) genes in male S2 cells. Genes > 3000 bp (n = 3152) were considered, the gene body (TSS + 1000 bp – TTS – 1000 bp) was scaled to 2000 bp and 500 bp up- and downstream of the genes were included.

Supplementary Figure 7 - Albig et al.

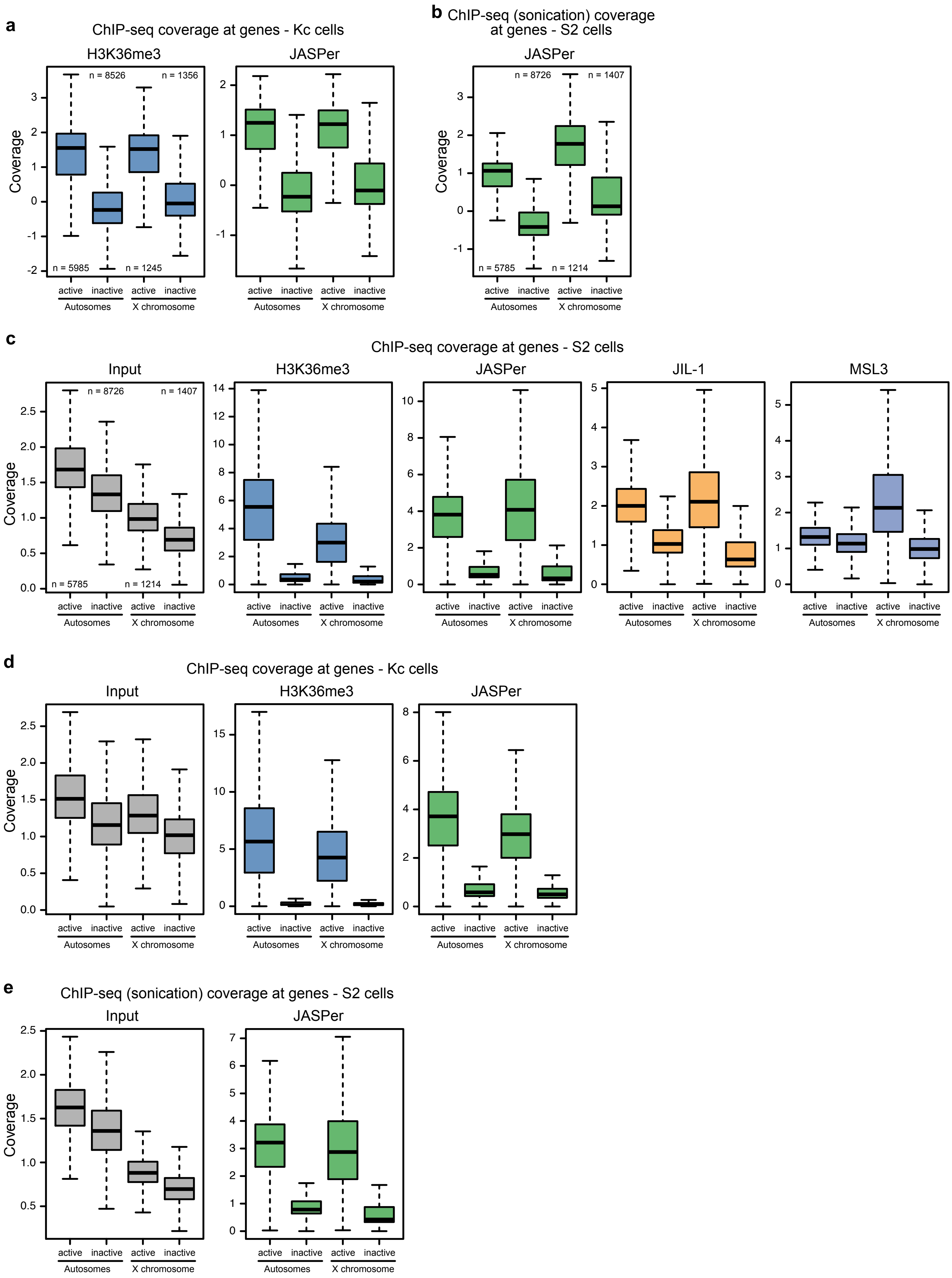

**Supplementary Figure 7. Coverage of X chromosome and autosomes in S2 and Kc cells for H3K36me3, JASPer, JIL-1 and MSL3 for comparison.**

(a) Box plot showing mean H3K36me3 (left panel) and JASPer (right panel) MNase ChIP-seq normalized coverage (n = 3 each) at active (tpm > 1) and inactive (tpm ≤ 1) genes on the autosomes (n = 5985 and n = 8526, respectively) and X chromosome (n = 1245 and n = 1356, respectively) in female Kc cells, as in Fig. 3d. Outliers are not shown.

(b) Box plot showing mean JASPer sonication ChIP-seq normalized coverage (n = 2) at active and inactive genes on the autosomes (n = 5785 and n = 8726, respectively) and X chromosome (n = 1214 and n = 1407, respectively) in male S2 cells, as in Fig. 3d. Outliers are not shown.

(c) Box plot showing mean Input (left panel, n = 7), H3K36me3 (second left panel, n = 4) and JASPer (second right panel, n = 4) and JIL-1 (right panel, n = 5) MNase ChIP-seq non-input normalized coverage at active (tpm > 1) and inactive (tpm ≤ 1) genes on the autosomes (n = 5985 and n = 8526, respectively) and X chromosome (n = 1245 and n = 1356, respectively) in male S2 cells. Outliers are not shown.

(d) Box plot showing mean Input (left panel), H3K36me3 (middle panel) and JASPer (right panel) MNase ChIP-seq non-input normalized coverage (n = 3 each) at active (tpm > 1) and inactive (tpm ≤ 1) genes on the autosomes (n = 5985 and n = 8526, respectively) and X chromosome (n = 1245 and n = 1356, respectively) in female Kc cells, as in (c). Outliers are not shown.

(e) Box plot showing mean Input (left panel) and JASPer (right panel) sonication ChIP-seq non-input normalized coverage (n = 2 each) at active (tpm > 1) and inactive (tpm ≤ 1) genes on the autosomes (n = 5785 and n = 8726, respectively) and X chromosome (n = 1214 and n = 1407, respectively) in male S2 cells, as in (c). Outliers are not shown.

Supplementary Figure 8 - Albig et al.

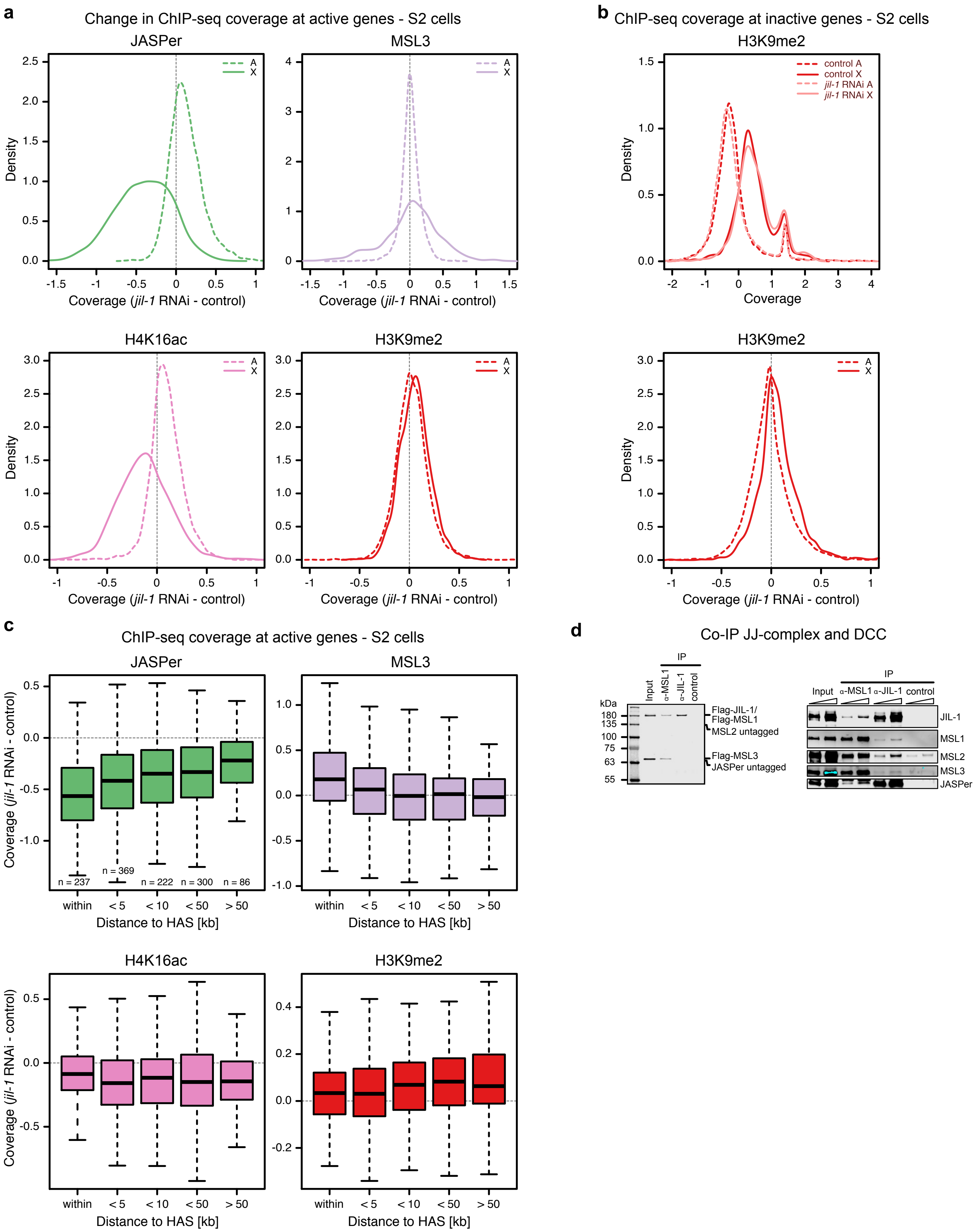

**Supplementary Figure 8. Changes in JASPer, MSL3, H4K16ac and H3K9me2 coverage at active genes on the X versus autosomes in S2 cells JIL-1 depletion.**

(a) Density plot showing difference of mean (n = 3 each, for MSL3 n = 2) spike-in ChIP-seq normalized coverage after *jil-1* RNAi treatment and control male S2 cells at active (tpm > 1) genes for JASPer (top left), MSL3 (top right), H4K16ac (bottom left) and H3K9me2 (bottom right). X chromosomal genes (n = 1214) are marked with solid line and autosomal genes (chromosomes 2L, 2R, 3L and 3R, n = 5785) with dashed line.

(b) Density plot showing mean (n = 3 each) spike-in ChIP-seq normalized coverage in control male S2 cells and after *jil-1* RNAi treatment at inactive (tpm ≤ 1) genes for H3K9me2 in upper panel. X chromosomal genes (n = 1407) are marked with solid line and autosomal genes (chromosomes 2L, 2R, 3L and 3R, n = 8726) with dashed line. In lower panel, difference of mean (n = 3 each) spike-in ChIP-seq normalized coverage after *jil-1* RNAi treatment and control male S2 cells at genes for H3K9me2.

(c) Box plot showing difference of normalized mean (n = 3 each, for MSL3 n = 2) spike-in ChIP-seq coverage after *jil-1* RNAi treatment and control male S2 cells at active (tpm > 1) X-linked genes for JASPer (top left), MSL3 (top right), H4K16ac (bottom left) and H3K9me2 (bottom right) binned according to their distance to HAS. The number of genes in each bin are indicated in the plot. Outliers are not shown.

(d) Western blot analysis using $\alpha$-FLAG antibody of co-IP experiments with extracts from Sf21 cells expressing wild type, untagged JASPer, FLAG-JIL-1, FLAG-MSL1, untagged MSL2 and FLAG-MSL3. Co-IP was performed with $\alpha$-MSL1 serum, $\alpha$-JIL-1 serum and non-specific serum as control. Left panel shows Western blot analysis using $\alpha$-FLAG antibody and right panel using protein specific antibodies.

Supplementary Figure 9 - Albig et al.

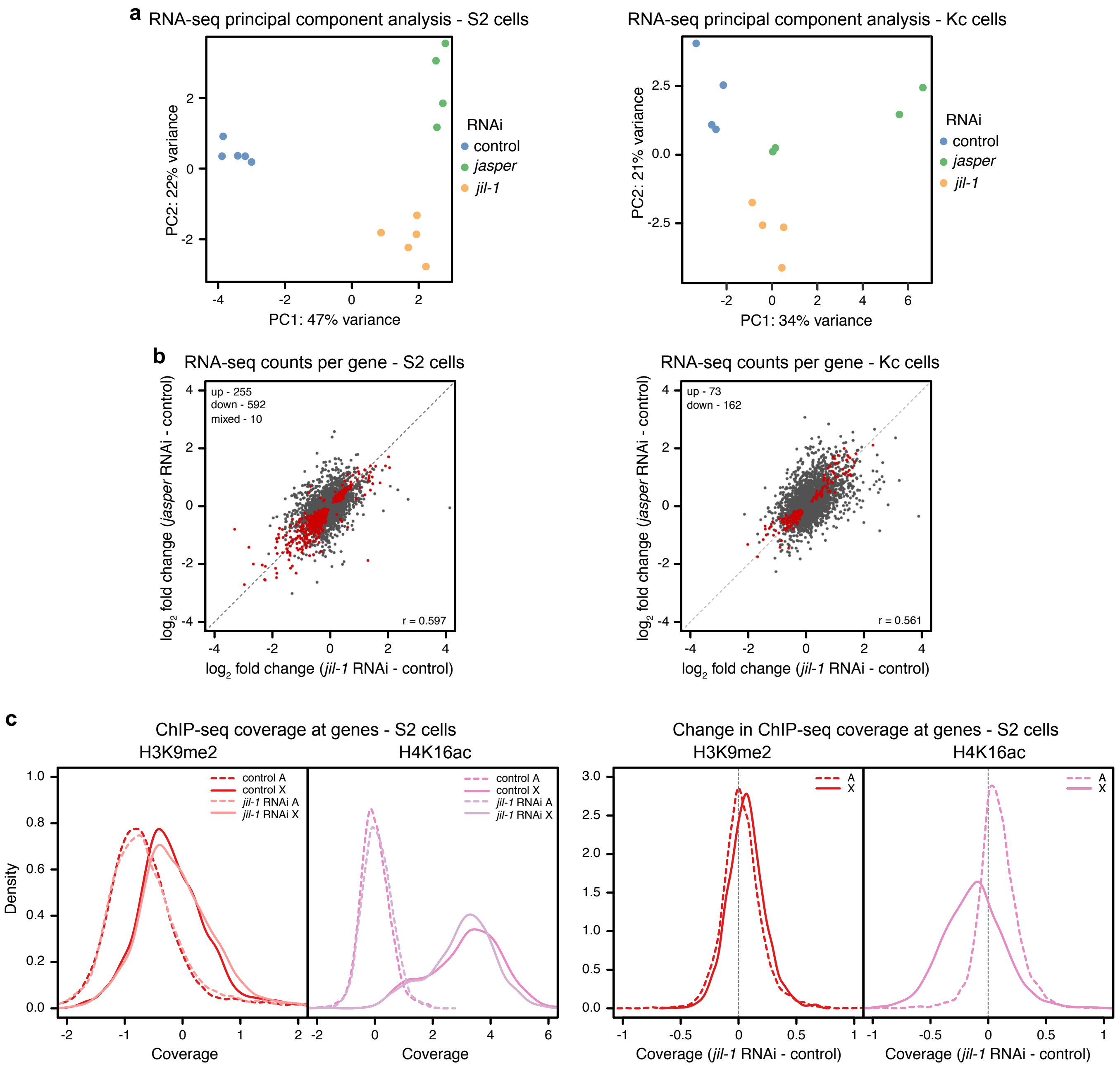

**Supplementary Figure 9. Gene expression changes after *jasper* and *jil-1* RNAi in cells and gene coverage of H3K9me2 and H4K16ac.**

(a) Scatter plot of PCA of RNA-seq counts after control (n = 5), *jasper* (n = 4) and *jil-1* RNAi (n = 5) at robustly detected genes (n = 8452) in male S2 cells showing PC2 against PC1, in left panel. Right panel, PCA of RNA-seq counts after control, *jasper* and *jil-1* RNAi (n = 4 each) at genes in female Kc cells (n = 8834).

(b) Scatter plot showing mean $\log_2$ fold-change RNA-seq counts upon *jasper* RNAi versus controls (n = 4) against *jil-1* RNAi versus controls (n = 5) at robustly detected genes in male S2 cells (n = 8452) with Pearson correlation coefficient r = 0.597 in left penal. Statistically significant differentially expressed genes between RNAi and control conditions (fdr < 0.05) common in both RNAi conditions are marked in red and the number of significant genes is indicated on the plot. Right panel, mean $\log_2$ fold-change of sRNA-seq counts upon *jasper* RNAi versus controls against *jil-1* RNAi versus controls (n = 4 each) at genes in female Kc cells (n = 8834) with Pearson correlation coefficient r = 0.561.

(c) Density plot showing mean (n = 3 each) spike-in ChIP-seq normalized coverage in control male S2 cells and after *jil-1* RNAi treatment at in RNA-seq robustly detected genes for H4K16ac (right) and H3K9me2 (left) in left panel. X chromosomal genes (n = 1509) are marked with solid line and autosomal genes (chromosomes 2L, 2R, 3L and 3R, n = 7144) with dashed line. In right panel, difference of mean (n = 3 each) spike-in ChIP-seq normalized coverage after *jil-1* RNAi treatment and control male S2 cells at genes for H4K16ac (right) and H3K9me2 (left).

Supplementary Figure 10 - Albig et al.

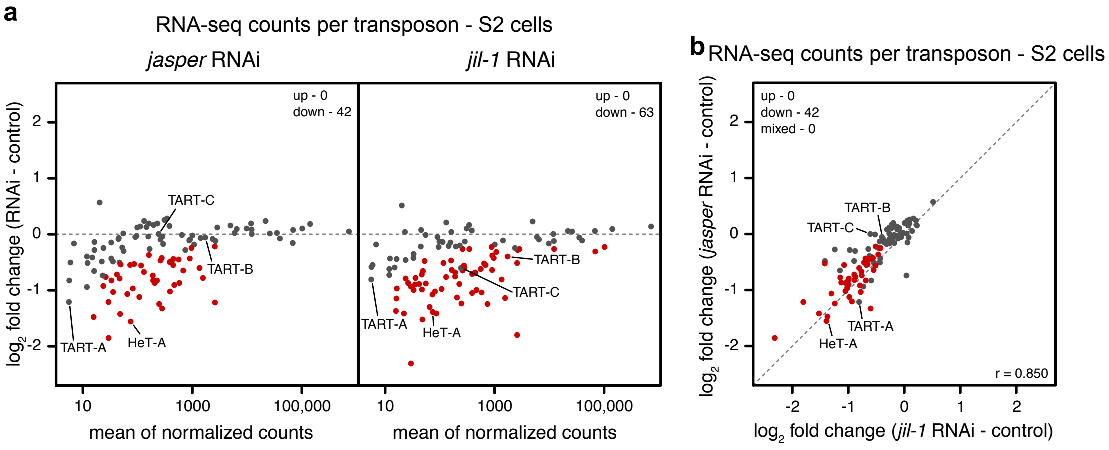

**Supplementary Figure 10. Expression changes of transposons after *jasper* and *jil-1* RNAi in cells.**

(a) Scatter plot showing mean $log_2$ fold-change of RNA-seq counts upon *jasper* RNAi versus controls (n = 4) and *jil-1* RNAi versus controls (n = 5) against mean RNA-seq counts at robustly detected transposons in male S2 cells (n = 111). Statistically significant differentially expressed transposons between RNAi and control conditions (fdr < 0.05) are marked in red and the number of significant genes is indicated on the plot. TEs of the HTT arrays are labeled.

(b) Scatter plot showing mean $log_2$ fold-change of RNA-seq counts upon *jasper* RNAi versus controls (n = 4) against *jil-1* RNAi versus controls (n = 5) at transposons in male S2 cells (n = 111), as in (a), with Pearson correlation coefficient r = 0.850, in left panel, in left panel. Statistically significant differentially expressed transposons between RNAi and control conditions (fdr < 0.05) common in both RNAi conditions are marked in red and the number of significant genes is indicated on the plot. TEs of the HTT arrays are labeled.

Supplementary Figure 11 - Albig et al.

**a**

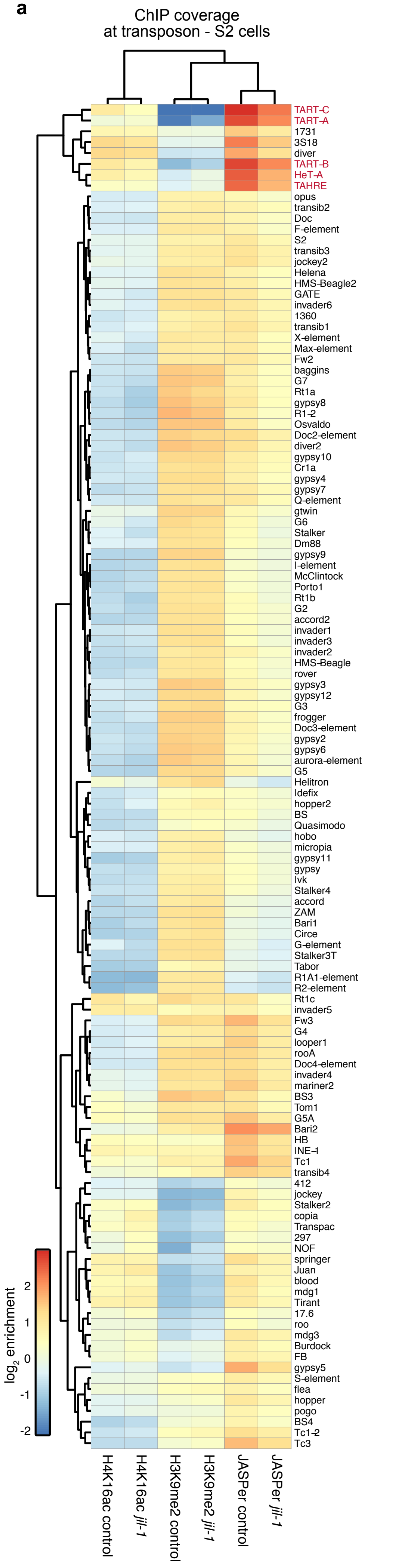

**Supplementary Figure 11. Transposons of the telomeric HTT arrays acquire H3K9me2 upon JIL-1 depletion in S2 cells.**

(a) Heatmap showing mean normalized $\log_2$ enrichment in H3K9me2 (n = 3), JASPer (n = 3) and H4K16ac (n = 3) MNase ChIP-seq at transposons (n = 124) in control male S2 cells and after *jil-1* RNAi. Transposons of the HTT array are marked in red.

Supplementary Figure 12 - Albig et al.

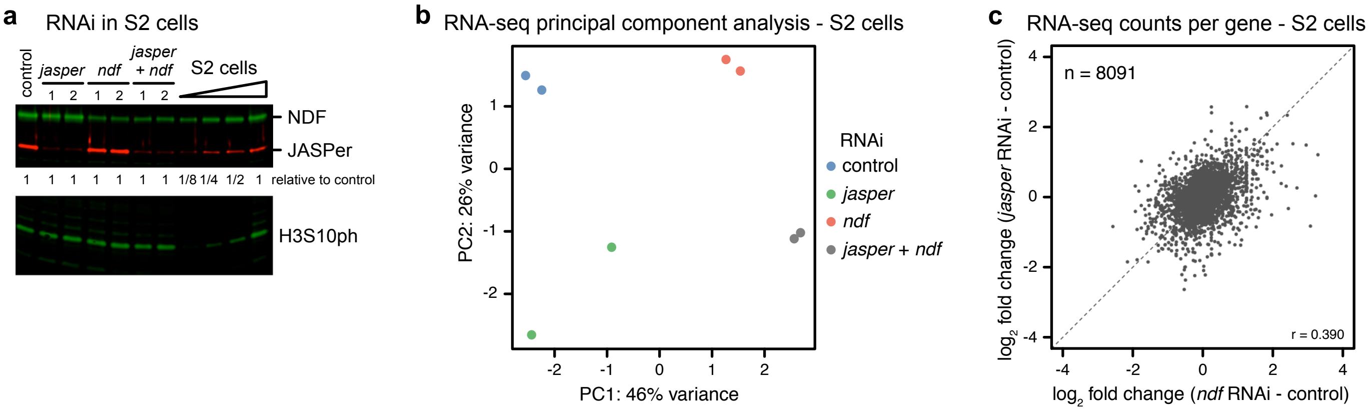

**Supplementary Figure 12. Double knock down of JASPer and dNDF has additive effects on gene expression.**

(a) Western blot analysis using $\alpha$-JASPer, $\alpha$-NDF and $\alpha$-H3S10ph antibodies of cell extracts from S2 cells after *jasper* and/or *ndf* or control RNAi treatment and dilution series of untreated S2 cell extract as reference. Ratio of extract loaded relative to control treatment according to cell number as indicated below.

(b) Scatter plot of PCA of RNA-seq counts after control, *jasper*, *ndf* and *jasper* + *ndf* RNAi (n = 2 each) at robustly detected genes in male S2 cells (n = 8091) showing PC2 against PC1.

(c) Scatter plot showing mean $\log_2$ fold-change of RNA-seq counts upon *jasper* RNAi versus controls against *ndf* RNAi versus controls (n = 2 each) at robustly detected genes in male S2 cells (n = 8091) with Pearson correlation coefficient r = 0.390.

**Supplementary Table 1. The JJ-complex interaction network.**

Table of complete IP-MS data reporting mean $\log_2$ fold enrichment in $\alpha$-JASPer IP (n = 6) versus control IP (n = 5) and associated p-values.
